# Membrane curvature sensing and stabilization by the autophagic LC3 lipidation machinery

**DOI:** 10.1101/2022.05.03.490522

**Authors:** Liv E. Jensen, Shanlin Rao, Martina Schuschnig, A. King Cada, Sascha Martens, Gerhard Hummer, James H. Hurley

## Abstract

How the highly curved phagophore membrane is stabilized during autophagy initiation is a major open question in autophagosome biogenesis. Here, we use *in vitro* reconstitution on membrane nanotubes and molecular dynamics simulations to investigate how core autophagy proteins in the LC3 lipidation cascade interact with curved membranes, providing insight into possible roles in regulating membrane shape during autophagosome biogenesis. ATG12–5-16L1 was up to 100-fold enriched on highly curved nanotubes relative to flat membranes. At high surface density, ATG12–5-16L1 binding increased the curvature of the nanotubes. While WIPI2 binding directs membrane recruitment, the amphipathic helix *α*2 of ATG16L1 is responsible for curvature sensitivity. Molecular dynamics simulations revealed that helix *α*2 of ATG16L1 inserts shallowly into the membrane, explaining its curvature-sensitive binding to the membrane. These observations show how the binding of the ATG12–5-16L1 complex to the early phagophore rim could stabilize membrane curvature and facilitate autophagosome growth.

## Introduction

Macroautophagy, hereafter autophagy, is the process in which cytosolic cargoes such as protein aggregates, damaged organelles, intracellular pathogens, or bulk cytoplasm are engulfed in a double-membrane vesicle and targeted to the lysosome for degradation^1^. Autophagic dysfunction results in defects in the clearance of aggregates and damaged organelles, contributing to human neurodegenerative diseases, including Parkinson’s Disease^2,3^. During cargo engulfment, the cup-shaped phagophore grows progressively larger^4^ and ultimately closes into the double-membraned autophagosome. The past few years have brought rapid progress in understanding how autophagy is initiated, cargo selected, and lipids are sourced and transferred for autophagosome expansion ^5–7^. In this context, the physical mechanism of membrane shaping and stabilization during autophagosome growth remains one of the most prominent open questions.

The formation and stabilization of the cup-shaped membrane during phagophore expansion is associated with energetic penalties and barriers^8–10^. Recent cryo-ET visualization of phagophores in yeast revealed that the membrane curvature at the phagophore rim approaches the maximum value possible given the thickness of the phospholipid bilayer^11^. The recruitment of membrane curvature inducing proteins to the phagophore rim is currently a leading model to explain how its high curvature is stabilized^9^. A number of core autophagy proteins have been shown to sense membrane curvature^12^. The PI3KC3-C1 subunit ATG14 senses membrane curvature through its C-terminal BATS domain^13–15^. Both the human phospholipid transporter ATG2A^9^ and the *Arabidopsis* ATG5 subunit of the ATG12–5-16L1 complex^16^ localize to toroidal structures that appear to correspond to the phagophore rim. ATG3^17^ and the ATG12–5-16L1 complex^18^, which are both involved in the conjugation of ATG8 proteins to membranes, preferentially bind to small liposomes through amphipathic helices at or near their N-termini. Thus, both cell imaging^16^ and *in vitro* binding^18^ data suggested that the ATG12–5-16L1 complex could stabilize the phagophore rim via a preference for high curvature membrane binding.

Covalent conjugation of the ATG8 family proteins LC3A-C, GABARAP, and GABARAPL1/2 to the membrane lipid phosphatidylethanolamine (PE), hereafter referred to as “LC3 lipidation”, contributes to phagophore expansion^19^ and the recruitment of cargo and other autophagy proteins^20,21^. LC3 lipidation proceeds through a cascade of enzymes analogous to the ubiquitin E1/E2/E3 ligase mechanism. The E1 ATG7 binds LC3, handing it off to the E2 ATG3, which works together with the E3 ATG12–5-16L1 to covalently conjugate LC3 to PE headgroups on the phagophore. The ATG12–5-16L1 is recruited to PI(3)P-positive autophagic membranes by the PROPPIN WIPI2 through its binding to a WIPI2 interacting region in ATG16L1^22,23^. These reactions have been reconstituted *in vitro* on sonicated liposomes^17,24^ and flat membranes^25^. In this study, we sought to systematically examine the impact of curvature on this machinery using a precisely tunable and quantitative *in vitro* system, and to gain a detailed molecular view of the membrane interactions with the help of molecular dynamics simulations.

Optical tweezers can be used pull membrane nanotubes from a giant unilamellar vesicle (GUV) and form a contiguous membrane system with regions of high and low curvature, enabling quantitation of the curvature sensitivity of proteins^26,27^ (Fig. 1a). This experimental setup can access nanoscale dimensions similar to the curvature of the phagophore rim (<30 nm^12^). The physics of membrane nanotubes pulled under force from an optical trap allows their radius to be measured by fluorescence microscopy even for dimensions far below the optical diffraction limit. Here, we used this approach to characterize the WIPI2 and ATG12–5-16L1 system and find that it is profoundly curvature sensitive.

**Figure 1.**
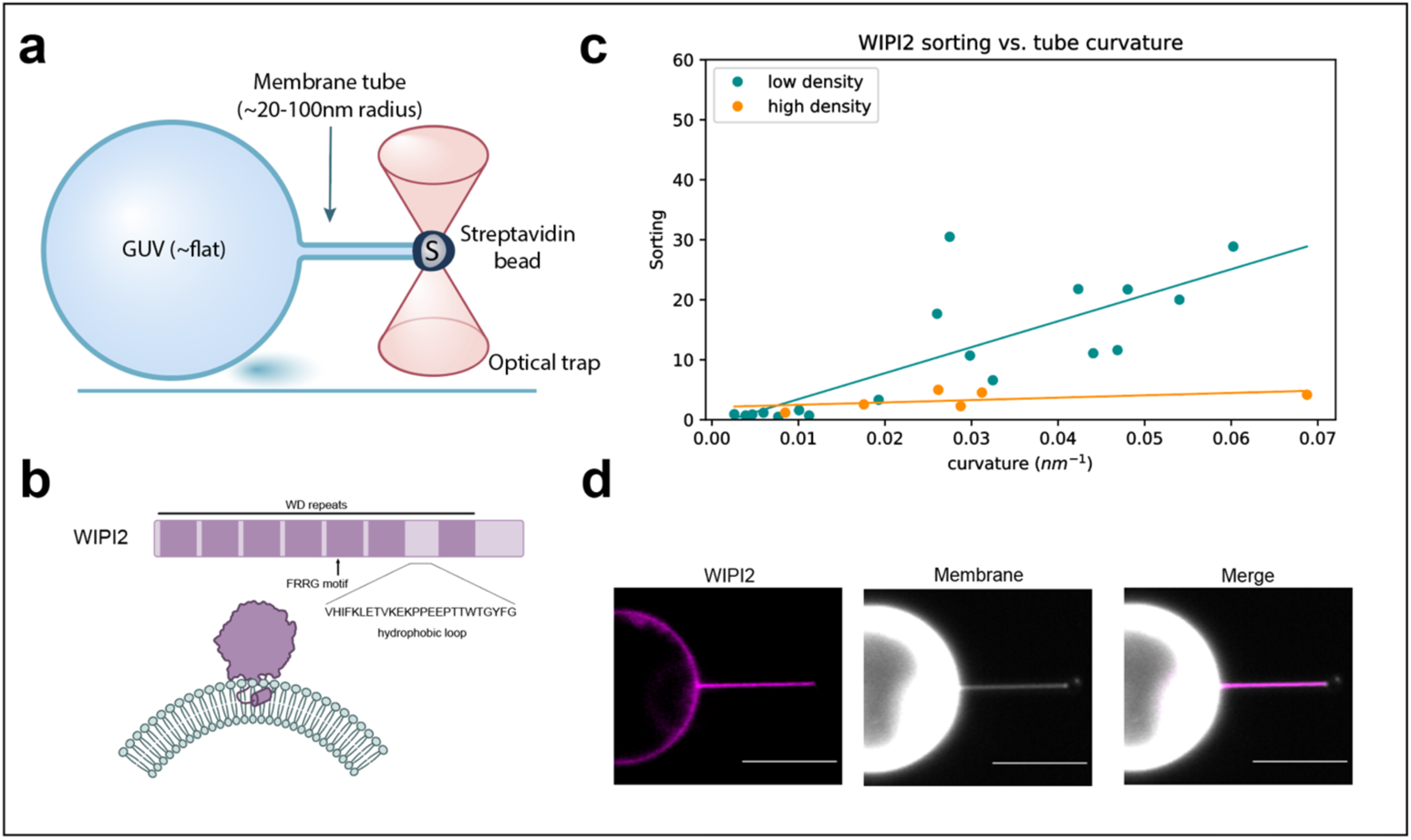
WIPI2 senses membrane curvature. (a) Schematic of GUV/optical trap setup for quantification of membrane curvature sensing. (b) Graphical representation of WIPI2 domain architecture and interaction with curved membrane. (c) Plot of WIPI2 sorting vs. membrane tube curvature at high (n=6) and low (n=18) surface densities of WIPI2 on the GUV surface. Each data point represents an individual membrane tube. (d) Representative confocal fluorescence microscopy images of WIPI2 localization on membrane tube and GUV surface. Scale bar: 10µm.

## Results

### WIPI2 interactions with membrane tubes

WIPI2 is responsible for the membrane recruitment of ATG12–5-16L1 in cells^22,23^, so we began by assessing how WIPI2 interacts with membranes of varying curvature. WIPI2 is a member of the PROPPIN family of proteins, which bind membranes through a hydrophobic loop^28^ together with an FRRG motif which specifically recognizes PI(3)P^28–31^ (Fig. 1b). We used the membrane nanotube assay to measure the curvature sensitivity of WIPI2 by visualizing WIPI2 localization to highly curved membrane tubes and the essentially flat GUV from which they were pulled (Fig. 1c). PI(3)P-positive GUVs were incubated with fluorescently labeled mCherry-WIPI2 and the curvature sensitivity of WIPI2 was assessed by its sorting ratio (*S*, the ratio of protein surface density on the tube to the protein density on the GUV surface^26^). At the limit of low density, the sorting ratio describes the tendency of membrane bound proteins to preferentially associate with the curved tube compared to the molecularly flat GUV surface.

Analysis of WIPI2 localization showed that it preferentially binds to membrane tubes compared to the GUV surface, and more strongly sorts onto narrow tubes compared to wide tubes. The sorting dependence on curvature for WIPI2 is most apparent on GUVs with a low protein surface density (ϕ_v_), reaching S_w_ = 14± 5 for tubes with radii between 20 nm and 40 nm (Fig. 1c, d). As expected, the sorting ratio of WIPI2 diminishes with increasing protein surface density on the GUV, which is attributable to approaching surface area saturation on the tube ^27,32^. A linear fit shows a monotonic increase in sorting with curvature, similar to proteins that are known to insert into lipid bilayers via amphipathic helices^33^.

To test whether the observed curvature-dependent sorting of WIPI2 is intrinsic to the protein itself and not due to enrichment of PI(3)P on the membrane tubes, we analyzed the localization of a fluorescently labeled PI(3)P probe, mCherry-FYVE. We found that, in contrast to WIPI2, mCherry-FYVE exhibited no enrichment on membrane tubes compared to the GUV surface (Extended Data Fig. 1a), indicating that sorting of WIPI2 is due to its inherent membrane curvature sensing activity.

### ATG12–5-16L1 has enhanced curvature sorting

Because both WIPI2 and ATG16L1 contain membrane binding motifs, and because WIPI2 is required to recruit ATG12–5-16L1 to flat membranes ^25^, we sought to understand the relative roles of WIPI2 and ATG16L1. We pulled membrane nanotubes from GUVs incubated with mCherry-WIPI2 and ATG12–5-16L1-GFP and analyzed their respective recruitment to the membrane tube (Fig. 2a, b, c). Just as for WIPI2, the sorting index *S* of ATG12–5-16L1 depends on protein surface density. Lower protein density on the GUV correlated with strong sorting of ATG12–5-16L1 onto membrane tubes (Fig. 2a). ATG12–5-16L1 was more strongly enriched on membrane tubes than WIPI2, with a sorting ratio S_E3_=63±35 for membrane tubes with radii between 20nm and 40nm, compared to a sorting ratio S_W_=15 ± 4 for WIPI2(Fig. 2b, d).

**Figure 2.**
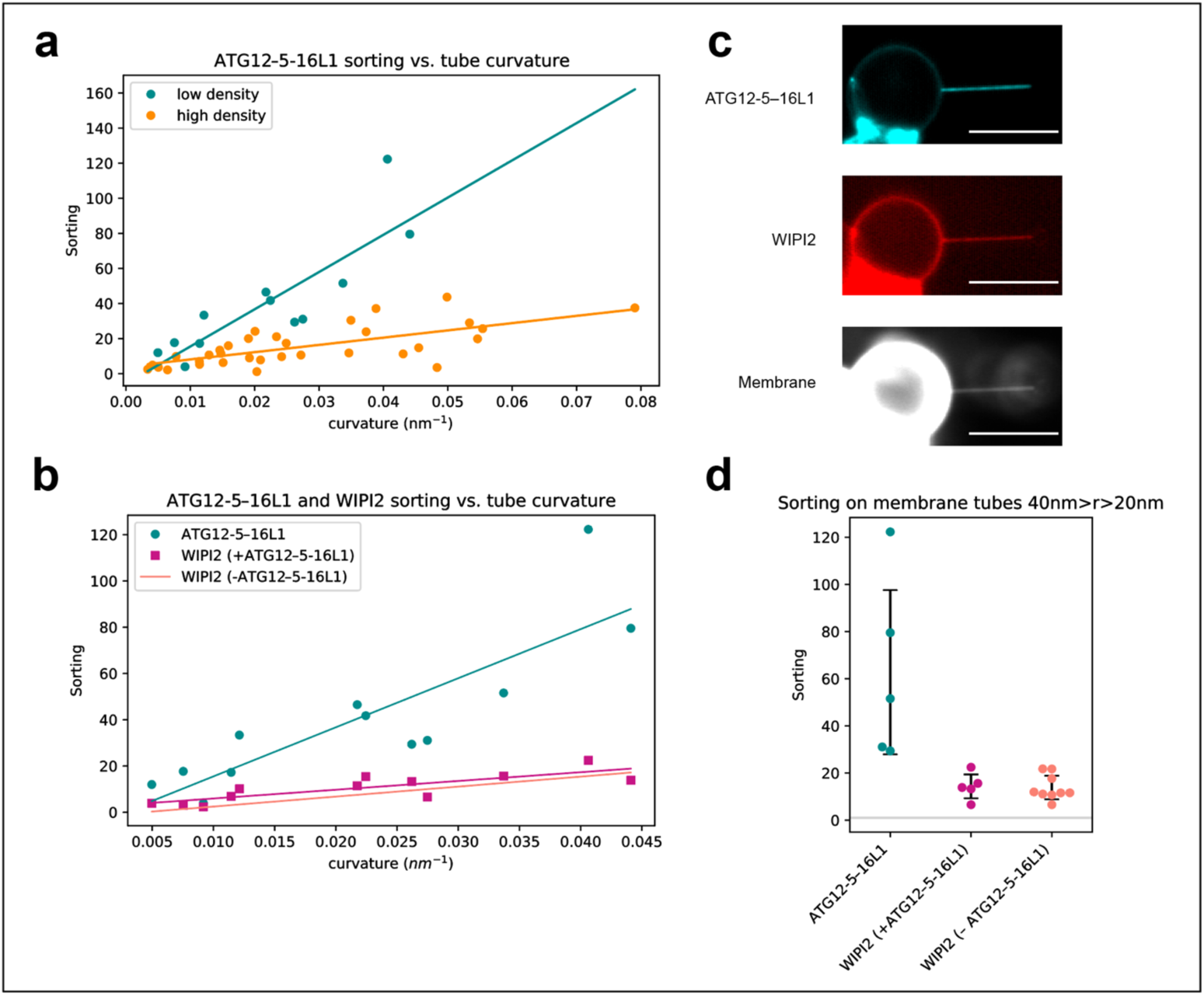
ATG12–5-16L1 has enhanced curvature sensitivity compared to WIPI2. (a) Quantification of ATG12–5-16L1 sorting onto membrane tubes at high (cyan, n=12) and low (orange, n=33) membrane surface densities of protein. Each data point represents an individual membrane tube. (b) Comparison of sorting dependence on curvature for ATG12–5-16L1 (cyan) and WIPI2 (magenta) with two-color fluorescence imaging compared to sorting of WIPI2 in the absence of ATG12–5-16L1 (salmon). Scale bar: 10µm. (c) Representative image of membrane tube enrichment of ATG12–5-16L1 and WIPI2 from (a) and (b). (d) Swarmplot depicting the sorting of ATG12–5-16L1 and WIPI2 (n=5), compared to WIPI2 in the absence of ATG12–5-16L1 (n=9) on membrane tubes with radii between 20and 40 nm, black bars indicate standard deviation, and horizontal grey line at S=1.

### ATG16L1 and WIPI2 curvature sensing is independent of ATG3

Having established the potent membrane curvature sensitivity of WIPI2 and ATG12–5-16L1, we next tested their curvature sensitivity in the presence of ATG3, the LC3 lipidation E2 enzyme. Previously identified as a membrane curvature sensor through an N-terminal amphipathic helix^17^, we hypothesized that ATG3 might be able to even further increase the curvature dependent sorting of WIPI2 or ATG12–5-16L1 through a direct interaction.

First, we tested the interaction between ATG3 and ATG12–5-16L1. ATTO565 labeled ATG3 was incubated with GUVs alone or in the presence of unlabeled WIPI2 and ATG12–5-16L1-GFP. ATG3 was only recruited to the GUV surface in the presence of ATG12–5-16L1, and co-partitioned with ATG12–5-16L1 onto membrane tubes (Fig. 3a). To assess the impact of ATG3 on the curvature sensitivity of ATG12–5-16L1 while controlling for potential differences in sorting induced by the fluorescent tags, we incubated GFP labeled ATG12–5-16L1 with unlabeled ATG3 and mCherry-WIPI2. We found that addition of ATG3 did not alter the partitioning of ATG12– 5-16L1 onto membrane tubes compared to ATG12–5-16L1 in the presence of WIPI2 alone (Fig. 3b). Despite the importance of its amphipathic helix for LC lipidation activity ^17^, ATG3 did not measurably augment the curvature sensitivity of WIPI2 and ATG12–5-16L1.

**Figure 3.**
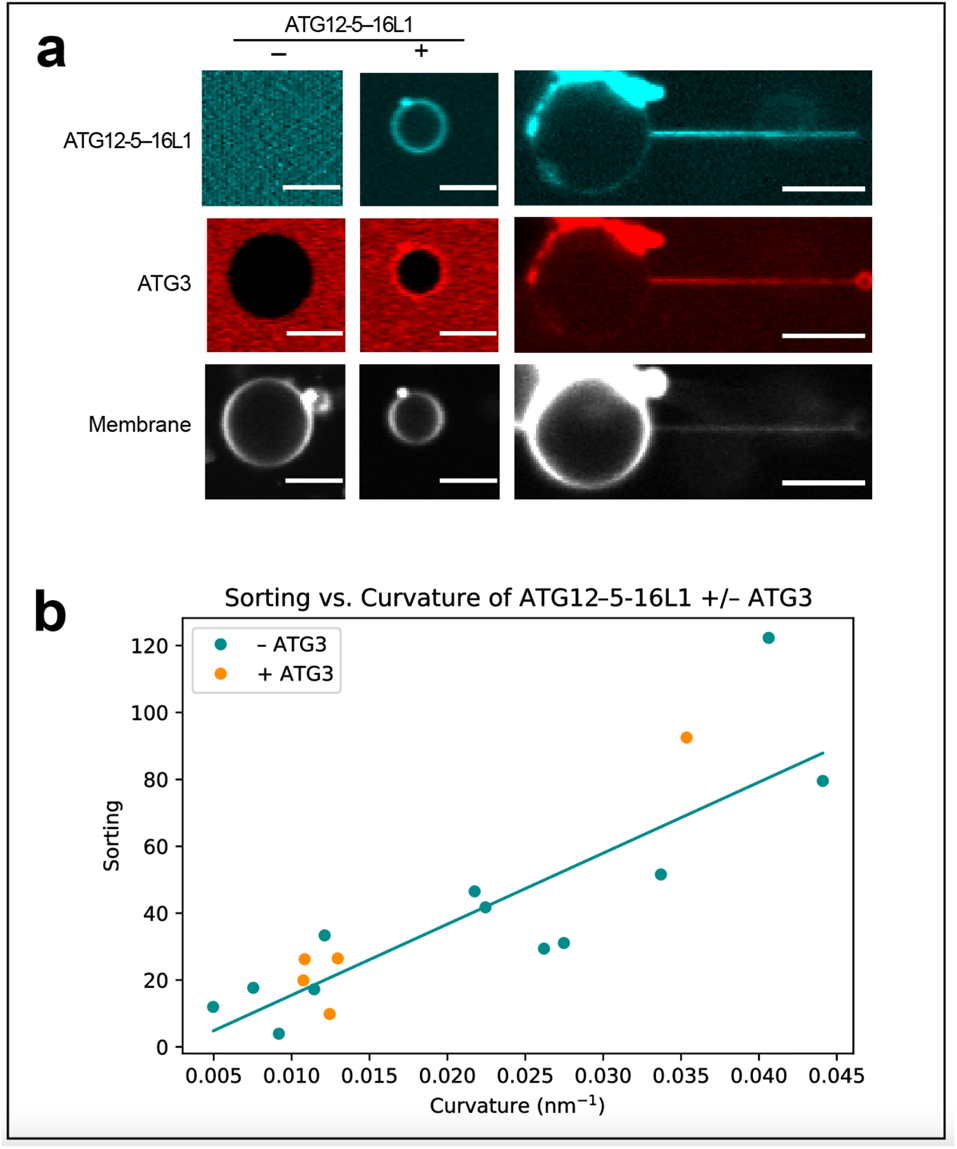
ATG12–5-16L1 senses curvature independently from ATG3. (a) ATG3 localization to GUVs and membrane tubes in the presence and absence of ATG12–5-16L1. Scale bar: 10µm. (b) Curvature dependence of ATG12–5-16L1 sorting on membrane tubes in the presence (n=5) and absence (n=12, from Fig. 2a) of ATG3. Each data point represents an individual membrane tube.

### ATG16L1 helix *α*2 is responsible for curvature sensitivity

We next sought to understand the mechanism for the potent curvature sensitivity of ATG12–5-16L1 complex. A previous report had characterized the membrane interactions of amphipathic helix *α*2, located near the N-terminus of ATG16L1^18^. On highly curved sonicated vesicles, the physiological requirement for WIPI2 membrane recruitment can be bypassed^25^. Helix *α*2 was shown to be essential for WIPI2-independent activity of ATG12–5-16L1 on sonicated liposomes, and essential for LC3 lipidation and autophagic flux in both bulk and selective autophagy in cells ^18^. We use the same FII mutation previously shown to block autophagy and LC3 lipidation in cells and assessed its impact on curvature sorting of ATG12–5-16L1.

A GFP-tagged version of the previously described FII mutant (F32A, I35A, I36A) of ATG16L1 was purified and incubated with GUVs in the presence of mCherry-WIPI2 (Fig. 4a, b). The FII mutation was sufficient to ablate the curvature sensitivity of ATG12–5-16L1 without impacting the sorting of WIPI2 onto membrane tubes (Fig. 4c, d). We conclude that the molecular basis for the amplified curvature-dependent sorting we observed for ATG16L1 depends on hydrophobic residues in the amphipathic helix *α*2 of ATG16L1.

**Figure 4.**
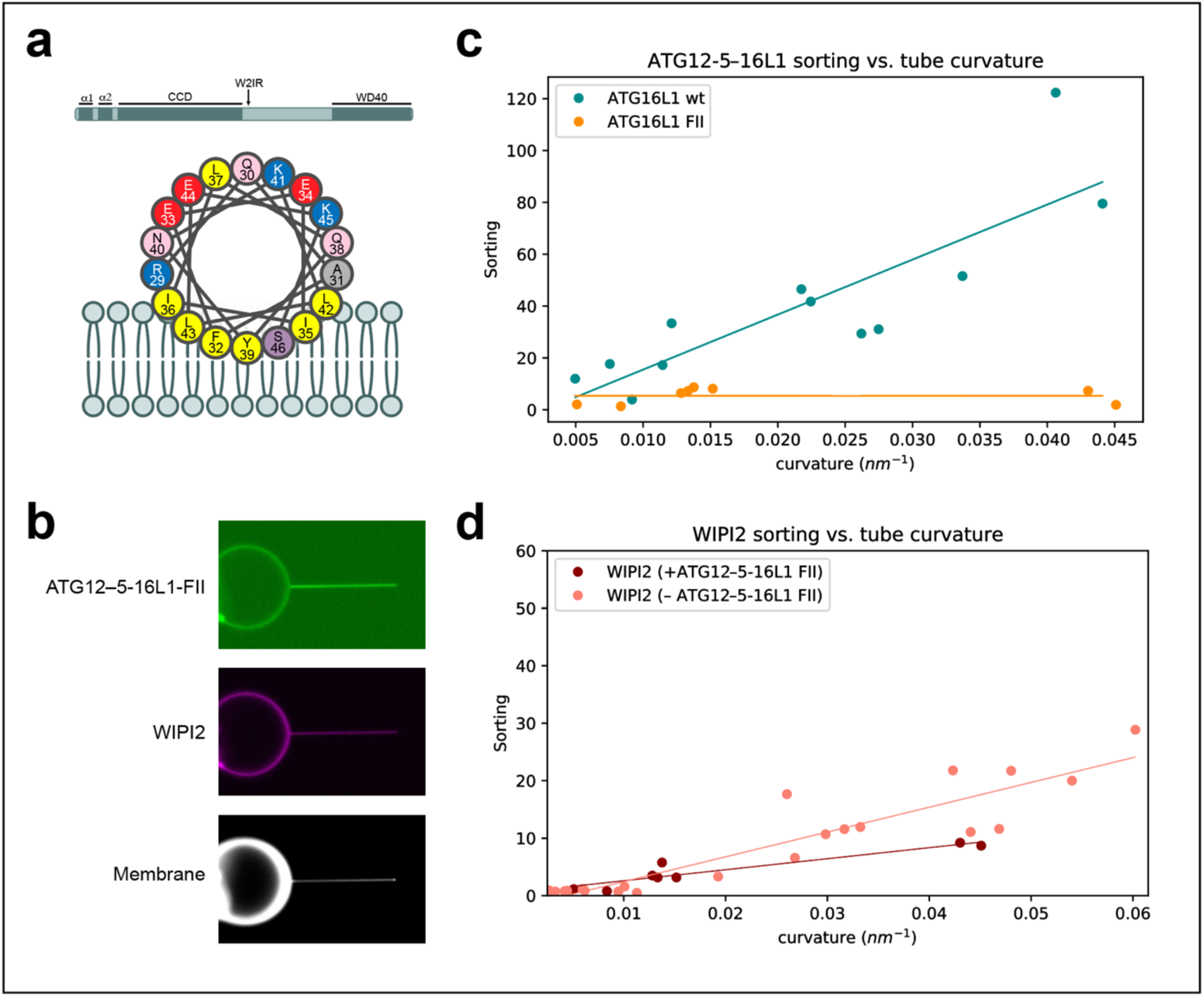
Mutagenesis impacts ATG12–5-16L1 curvature sensitivity. (a) Graphical representation of the domain architecture of ATG16L1 and of the *α*2 membrane binding amphipathic helix. (b) Representative fluorescence microscopy image of E3-FII and WIPI2 localization on GUV and membrane tube surface. (c) Sorting vs. curvature plot for ATG12–5-16L1 wt (n=12, from Fig. 2a) compared to FII mutant (n=8). (d) Sorting vs. curvature plot for WIPI2 alone (n=18, from Fig. 1c) or in the presence of ATG12–5-16L1 FII mutant (n=8).

### The ATG12–5-16L1 complex induces membrane curvature at high surface density

Thermodynamic principles of curvature sensing proteins dictate that at sufficiently high protein density, a curvature sensor becomes a curvature inducer^32^. Therefore, we wondered if ATG12–5-16L1 could induce membrane curvature at high surface densities at a physiologically plausible bulk concentration (100 nM). We analyzed time course images from nanotubes incubated with mCherry-WIPI2 and ATG12–5-16L1-GFP (Fig. 2), assessing how membrane tube radius changed with increasing protein surface density on the membrane tube. In these experiments, protein is preincubated with GUVs before the membrane tube is formed; after pulling the membrane tube, protein diffuses onto the tube from the GUV until it reaches its equilibrium sorting value. Taking advantage of the changing protein surface density, we measured the tube radius before and after protein enrichment on the membrane tube. Enrichment of ATG12–5-16L1 on membrane tubes consistently correlated with a concomitant decrease in tube radius (Fig. 5a). Visualizing a single representative tube over the course of several minutes shows that protein enrichment on the tube from ~600 to ~1700 dimers/µm^2^ induces an increase in curvature from 50nm to 25nm radius, as calculated from the decrease in the fluorescence signal from the membrane label in the tube (Fig. 5b, c). A decrease in fluorescence was not observed on tubes with already high curvature (Fig. 5a).

**Figure 5.**
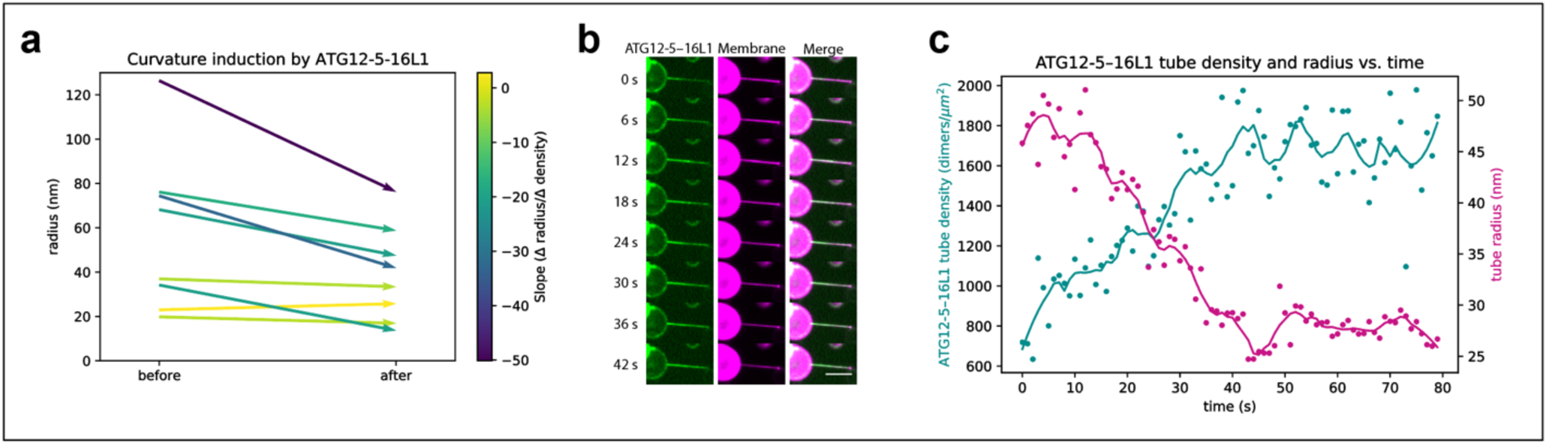
Curvature induction by ATG12–5-16L1. (a) initial and final tube radii after the accumulation of ATG12–5-16L1 onto the membrane tube. Vectors color coded for slope of the change in protein density over the change in tube radius, where darker colors indicate a steeper negative slope. (b) Montage of a single representative membrane tube showing ATG12–5-16L1 localization relative to the tube and membrane tube intensity over time. Snapshots of a continuous acquisition every 6s, showing ATG16L1-GFP (green) and ATTO647N-DOPE membrane (magenta). (c) Quantification of protein binding and membrane tube radius from timecourse in (b). Surface density of ATG16L1 dimers on the membrane tube (cyan) and membrane tube radius (magenta) plotted against time (s) for an 80 second timecourse.

### Molecular dynamics simulations of curved membrane binding

To clarify the mechanism of membrane association and curvature sensing by ATG16L1, we performed molecular dynamics simulations. Structural work has shown how the N-terminus of ATG16L1 interacts with the other subunits of the ATG12–5-16L1 complex via helix α1 ^34^. In atomistic molecular dynamics simulations of helix α2 of ATG16L1 as bound to ATG12–5 near negatively charged membranes (Fig. 6a), membrane association of ATG16L1 occurred within the first few nanoseconds of each 1 µs replicate. A stable interaction interface was maintained between α1 and ATG5 (Extended Data Fig. 2a). Helix α2 exhibited considerable flexibility relative to α1, with swinging and rotation about the hinge region around Gln30 and Ala31. This led to spontaneous reorientation (by up to ~190° relative to the initial conformation; Extended Data Fig. 2b) of the hydrophobic face of α2, with the side chains of Phe32 and Ile36 brought to the protein-membrane interface (Fig. 6b) in two of five replicates (two of ten ATG16L1 molecules simulated). These observations are consistent with membrane binding by the N-terminal region of ATG16L1 via exposure and insertion of hydrophobic side chains of α2, although membrane insertion events did not occur on the timescale of these simulations.

**Figure 6.**
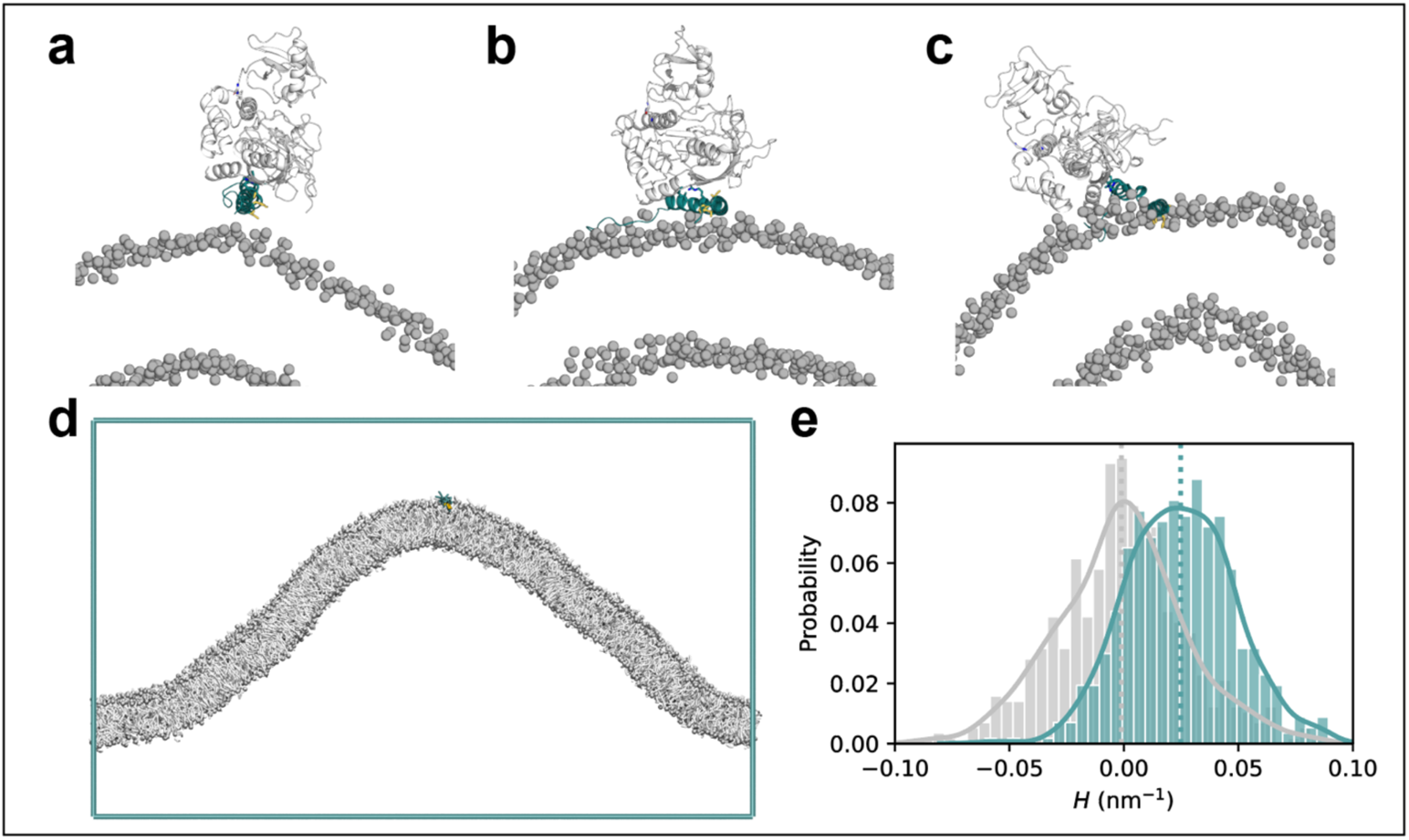
ATG16L1 helix 2 shows preference for positive curvature in molecular dynamics simulations. (a) Initial structure (PDB ID: 4NAW) of ATG16L1 helix α2 (cyan) in complex with ATG12-5 (white) near the membrane. The hydrophobic-face residues Phe32, Ile35, and Ile36 are highlighted as yellow sticks. Phosphorus atoms of lipid headgroups are shown as grey spheres. (b) Snapshot of ATG1612–5-L1 interacting with the membrane surface during one (1 µs) simulation replicate, with rotation of the hydrophobic face by ~180°. (c) End frame of a 300 ns replicate with helix α2 embedded into the membrane. (d) End frame of a 50 µs coarse-grained curvature sampling simulation replicate with ATG16L1 helix <u>α</u>2 (residues 26-45; located at the positively curved cusp of the membrane fold). (e) Probability histogram of local mean curvature values sampled at the center of mass of helix <u>α</u>2 (cyan) and at random lipid phosphate positions (grey) during the same simulations, with data collected over six independent 50 µs replicates, after an equilibration period of 5 µs upon each membrane binding event.

To examine the membrane binding configuration of ATG12–5-16L1 brought about by rotation of α2, a separate set of simulations was initiated using a remodeled structure of ATG16L1. Here, the hydrophobic face of α2 was embedded into the membrane prior to simulation (Fig. 6c). Over a total simulation time of ~1.8 µs across six independent copies of the molecule, the side chains of Phe32, Ile35, and Ile36 remained stably embedded within the membrane (Extended Data Fig. 2c), whilst α1 mediated contact with ATG12–5 above the membrane.

The curvature preference of helix *α*2 was assessed using coarse-grained molecular dynamics simulations of the isolated ATG16L1 construct (residues 26-45) on buckled membranes ^35^ (Fig. 6d). In these longer (50 µs) simulations, multiple membrane insertion and dissociation events were observed on a microsecond timescale. During its periods of membrane association, α2 diffused within the buckled membrane and showed a strong preference for regions with high positive local curvature. Sampling at the center of mass of α2 (and allowing for an equilibration period for the helix to reach its preferred membrane regions), the distribution of local mean curvature *H* of the membrane profile showed a clear shift towards positive curvature relative to random samples of lipid headgroup positions, with a mean value of *H*~0.025 nm^-1^ (Fig. 6e).

Residues 272-296 within the 6CD loop of WIPI2d^23^ were identified to constitute a candidate curvature-sensing element based on secondary structure prediction^36^, analysis of physicochemical properties^37^, and atomistic molecular dynamics simulations. The prediction of two short amphipathic α-helices, respectively consisting of residues 272-284 and 290-296 (Extended Data Fig. 2d), was corroborated by observations of spontaneous helix formation and membrane interaction in these regions during atomistic molecular dynamics simulations of WIPI2d near the membrane, initially with an unstructured 6CD loop. In further simulations of the protein with the two predicted helices pre-modeled into the structure, spontaneous membrane insertion of residues ~271-300 was observed within 1.5 µs in one replicate (Extended Data Fig. 2e). The two putative PI(3)P binding sites^28,30,38^ of WIPI2d, formed by blades 5 and 6 of the β-propeller, respectively, were also optimally positioned to bind PI(3)P in such a configuration (Extended Data. 2e). These observations agree with previous studies suggesting amphipathic helix formation and membrane insertion of the 6CD region in the orthologous Atg18 protein^39,40^. Coarse-grained curvature-sampling simulations of WIPI2d 6CD residues 263-300 on buckled membranes yielded a mean local curvature of *H*~0.015 nm^-1^ at the center of mass of the construct (Extended Data. 2f), supporting its role as a curvature sensor, albeit with weaker curvature sensitivity (and stronger membrane binding) compared with helix α2 of ATG16L1 (Fig. 6e).

Previous work modeling membrane association of membrane curvature generating proteins has highlighted the depth of helix insertion into the membrane as a key regulator of the magnitude of curvature sensitivity ^41^. We plotted the insertion depth of all-atom trajectories of WIPI2 loop and ATG16L1 *α*2 helix membrane insertion. We found that α2 of ATG16L1 interacts with the membrane at the level of the phosphate groups, whereas the WIPI2d 6CD loop inserts deeper into the membrane by ~5%-20% of the thickness of the monolayer (Extended Data. 3). Consistent with the membrane nanotube enrichment results, this finding supports a role for WIPI2 in strongly and stably associating with the membrane, whereas the ATG16L1 shallow membrane insertion is responsible for weaker binding with heightened membrane curvature sensitivity.

## Discussion

This study set out, in the first instance, to determine if ATG12–5-16L1 bound to membranes in a curvature-dependent manner in the physiologically relevant setting of WIPI2-driven recruitment. We found that WIPI2 is itself curvature-sensitive, but ATG12–5-16L1, following its recruitment by WIPI2, is even more sensitive. The sorting index for WIPI2-recruited ATG12–5-16L1 reaches a value of ~100 also seen for dedicated curvature sensors such as amphiphysin^42^. MD simulations show that WIPI2 inserts substantially into the hydrocarbon core of the membrane, while helix *α*2 of ATG16L1 inserts more shallowly. Shallow membrane binding by amphipathic helices has been shown to correlate with high curvature sensitivity^41^. Thus, the role of WIPI2 in this system is to drive membrane recruitment by binding tightly to PI(3)P and inserting deeply into the membrane, while ATG16L1 itself interacts weakly but in highly curvature dependent manner.

As noted previously^26^, curvature dependence is stronger at lower surface density. This likely represents the onset of saturation of the binding capacity of the membrane tube. It is also consistent with a model for curvature sensing in which self-association of ATG12–5-16L1 complexes with one another is neither required nor favorable. Curvature induction requires higher protein densities, at which the differential sorting of ATG12–5-16L1 to regions of high curvature is less evident.

There are some differences in the reported observations of ATG12–5-16L1 localization in cells between species. It is worth noting that ATG16L1 has almost negligible sequence similarity with yeast Atg16. The properties of yeast Atg16, which has been reported to tether^43^ and deform^44^ membranes, are therefore likely to differ from mammalian ATG16L1. In three-dimensional imaging of *Arabidopsis* cells, ATG5 is strikingly localized to toroidal zones that appear to correspond to the highly curved phagophore rim^16^. In mammalian cells, however, two-dimensional EM imaging of thin sections in mouse embryonic stem cells suggested that ATG16L1 is uniformly localized on phagophores^45^. We speculate that under conditions where ATG16L1 is strongly driving phagophore rim stabilization, higher surface densities are required, and therefore differential localization to the rim will be difficult to observe. This may be the case for the mouse ES cells as compared to *Arabidopsis* root epidermal cells in these studies. Moreover, theoretical considerations suggest rim stabilization is likely to be most important in the initial formation of the cup-shaped membrane. The apparent high density and uniform distribution of ATG12–5-16L1 across the phagophore in mouse cells^45^ likely serves the canonical function of the complex in LC3 lipidation at during the later stage of phagophore expansion, distinct from its early role in curvature stabilization.

WIPI2 and ATG12–5-16L1 are exemplars of curvature-sensitive autophagy proteins whose curvature preference is strong enough that it could, in principle, contribute to stabilizing phagophore rim curvature. They are unlikely to be the only such proteins. Indeed, the binding and/or activity of mammalian PI3KC3-C1^15^ and ATG3^17^ have been shown to be curvature sensitive *in vitro*. We found that ATG3 does not further increase the curvature sensitivity of ATG12–5-16L1, however, this is likely to reflect saturation of the already very high sorting index for ATG12–5-16L1, making it difficult to measure any further incremental increase. Indeed, saturation of the measurement may have led us to underestimate the true curvature dependence of ATG12–5-16L1 itself. It seems reasonable that both PI3KC3-C1 and ATG3 could contribute to phagophore rim stabilization. *In vitro* data are still lacking for ATG2A, but its apparent phagophore rim localization^9^ makes ATG2A an intriguing candidate. Under curvature inducing conditions, it might be expected that the rim would be saturated and broader localization of ATG2A observed. We therefore speculate that ATG2A is rim localized in order to deliver phospholipids to the rim for growth, as opposed to rim stabilization. At any rate, further biophysical and cell biological characterization of the curvature dependence of PI3KC3 and ATG2A, and comparison to the ATG12–5-16L1 data will be called for.

The data reported here help explain the previous report that ATG16L1 *α*2 is essential for autophagosome formation^18^. In this report, it was shown that recruitment of ATG16L1 to WIPI2 puncta, which mark sites of autophagy initiation, is unimpaired. Similarly, we found that GUV recruitment of the ATG12–5-16L1 FII mutant, which blocks membrane recruitment by *α*2, is unimpaired, because WIPI2 binding through the WIPI2-interacting region of ATG16L1 drives this process. The *α*2 mutant leaves intact the ATG12-5 unit, responsible for the recruitment of ATG3 and in turn, for LC3 lipidation. The most parsimonious explanation for the strong autophagic loss of function phenotype of ATG12–5-16L1 FII^18^ is that there is a loss of early phagophore rim stabilization.

## Data Availability Statement

Imaging data from membrane tube experiments has been uploaded at Zenodo (<u>10.5281/zenodo.6508734</u>). Molecular dynamics data has also been uploaded at Zenodo (<u>10.5281/zenodo.6513345</u>).

## Code Availability Statement

Code for image analysis has been uploaded at GitHub (https://github.com/livjensen7/Jensen_etal_2022)

## Methods

### Protein Expression and Purification

ATG12–5-16L1-GFP constructs (Addgene_169077) were expressed and purified from SF9 cells as described^25^ (dx.doi.org/10.17504/protocols.io.br6qm9dw). Briefly, cells were resuspended in lysis buffer (50mM HEPES pH7.5, 300mM NaCl, 2mM TCEP, + Complete protease inhibitor (Roche)), and lysed by sonication. Lysate was clarified by centrifugation, and the soluble fraction applied to a streptactin Sepharose column (Cytiva). Upon elution from the strep column with 5mM desthiobiotin (Sigma), fractions were concentrated by filter centrifugation, and purified by gel filtration over a Superose 6 column (Cytiva). mCherry-WIPI2d (Addgene_178912) was expressed and purified from suspension HEK GnTI cells^46^ (dx.doi.org/10.17504/protocols.io.bvjnn4me). Cells were resuspended in lysis buffer (50mM HEPES pH7.5, 200mM MgCl_2_, 10% glycerol, 1% Triton X-100, 1mM TCEP, +Complete protease inhibitor), and lysed by gentle rocking at 4°C for 30 minutes. Lysate was clarified by centrifugation, applied to a streptactin Sepharose column, and eluted with buffer containing 10mM desthiobiotin. GST-mCherry-FYVE (https://www.protocols.io/view/expression-and-purification-protocol-of-gst-mch-fy-b8k5ruy6) and His-TEV-ATG3 (Addgene_169079, https://www.protocols.io/view/expression-and-purification-protocol-of-homo-sapie-b8k2ruye) were expressed and purified from e. Coli BL21(DE3) culture.

### Protein Labeling

ATG3 was labeled with ATTO 565 NHS ester (ATTO-TEC). Briefly, 40µM ATG3 was mixed with 80µM ATTO 565 NHS ester in 50mM HEPES pH8.0, 150mM NaCl, 2mM TCEP. The reaction was carried out for 1 hour at room temperature, and buffer exchanged over a G-25 desalting column (Cytiva) into 50mM Tris, pH 8.0, 150mM NaCl, 2mM TCEP to quench the reaction and remove any unconjugated dye. Labeling efficiency was assessed by the ratio of absorbance at 280 and 564 nm, correcting for dye absorbance at 280nm, using a Nanodrop spectrophotometer. Protocol submitted to protocols.io.

### GUV preparation

GUVs were prepared by polyvinyl alcohol (PVA) assisted swelling. Briefly, 100ul of 5% PVA was spotted onto a glass coverslip and dried at 50°C. 50nMol of lipids dissolved in chloroform were mixed (mol percent: 70% DOPC, 20% DOPE, 5% DOPS, 5% DO-PI(3)P, 0.3% ATTO647N-DOPE, 0.01% PEG2000-biotin-DSPE) and dried on the PVA layer overnight in a vacuum desiccator. GUVs were swelled for 30-60 minutes at room temperature in 100ul of sucrose solution slightly hypotonic to imaging buffer (320mOsm) as determined by freezing point depression osmometer (Osmette III, Precision Systems). (https://www.protocols.io/view/guv-preparation-b8kzrux6)

### Membrane Tube Assay

Proteins were mixed with fluorescently labeled GUVs and then added to a microscope chamber that had been passivated with 1mg/ml BSA in imaging buffer and subsequently rinsed with imaging buffer (20mM Tris, pH 8.0, 150mM NaCl, 2mM MgCl_2_, 2mM TCEP). GUVs were allowed to settle before adding streptavidin coated silica beads (Spherotech) that had been diluted 1:1000 in imaging buffer. Using an optical trap, the bead was brought into contact with the biotinylated GUV surface and retracted to form a membrane tube. Protein bound to the GUV and tube membrane was monitored by confocal fluorescence imaging. (Protocol submitted at protocols.io)

### Imaging and Image Analysis

Imaging of the membrane tubes was performed on a Nikon Ti-Eclipse microscope with Nikon A1 confocal unit, modified with an optical trap and micromanipulators^47,48^, using a Plan Apochromat 60X/1.20 water immersion objective (Nikon). A complete image dataset for this paper can be found at (<u>10.5281/zenodo.6508734</u>). Image files were processed in ImageJ to generate regions of interest (ROIs) containing segments of the membrane tube or of the GUV surface. ROIs were combined and analyzed using custom Python scripts (https://github.com/livjensen7/Jensen_etal_2022). Briefly, the ROIs were segmented based on the intensity of the membrane dye channel with an Otsu threshold for local maxima, and protein binding signal was quantified as the average value in the protein label channel masked with the membrane channel segmentation. Background was calculated as the average value of pixels not in the membrane channel mask and subtracted from the signal as calculated above. Protein enrichment on membrane tubes (S) was calculated as a ratio of protein intensity on the tube to protein intensity on the GUV surface, normalized for membrane intensity:

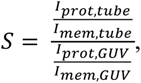

where *I_prot_* and *I_mem_* are the average intensity values from the ROI for the protein or membrane fluorescence channel, on the tube or GUV surface. (Protocol submitted at protocols.io).

### Tube Radius Calculation

Tube radii were calculated using a previously described method,^49^ in which the ratio of membrane dye fluorescence in the tube ROI to that in the GUV ROI is multiplied by an experimentally derived calibration constant (*k_tub_*):

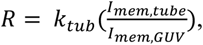

where *R* is the tube radius, and *I_mem,tube_* and *I_mem,GUV_* are the average intensity values of the membrane dye from the tube or GUV ROI.

To determine the value of *k_tub_*, tubes were pulled from GUVs held on an aspiration pipette, and the membrane tension varied by changing the aspiration force with a microfluidic controller (MFCS-EZ, Fluigent) to generate tubes of varying radii. Membrane tension (σ) was determined from aspiration pressure and microscopy images as:

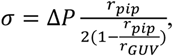

where Δ*P* is the difference in aspiration pressure from baseline, and *r_pip_* and *r_GUV_* are the radii of the aspiration pipette and the GUV, respectively.

Tube radius was subsequently calculated from membrane tension and force (*F)* felt by the bead in the optical trap as:

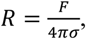

and plotted against the ratio of tube to GUV membrane fluorescence (Extended Data. 1b). Then, *k_tub_* was extracted from the slope of a linear least squares regression. (Protocol submitted at protocols.io)

### Surface Density Calculation

Protein density on the GUV surface was calculated by drawing a relationship between known concentrations of protein and membrane label in bulk solution and on the GUV surface^49^. First, standard curves of GFP and DOPE-ATTO488 were generated from a serial dilution in imaging buffer + 0.1% Triton X-100. The ratio of the slopes of the linear fits was used to correct for differences in optical properties of the different fluorophores. Then, the fluorescence intensity of GUVs containing a range of known percentages of dye-conjugated lipid (0.01-1 mol%) was calculated and plotted against surface density of the dye-conjugated lipid (assuming 0.7nm^2^/lipid) to give a relationship between intensity and GUV surface density. This was corrected by the ratio of intensities of the fluorophores in bulk solution. (Protocol submitted at protocols.io).

### Molecular dynamics simulations

Molecular dynamics simulations were performed with GROMACS 2020^50^, using the CHARMM36m force field^51^ for all-atom simulations and the MARTINI 3 force field^52^ and Gō-MARTINI model^53^ for coarse-grained systems.

Atomistic models of the ATG12–5-16L1 and WIPI2d-ATG16L1 complexes were based on crystal structures with PDB IDs 4NAW^54^ and 7MU2^23^, respectively. The ATG16L1 N-terminal helix in the former complex was replaced by a more complete structure (PDB ID: 4TQ0^55^) and residues 1-9 were added using the DEMO server^56^ to give a model of residues 1-50. A second model of the same region was generated in PyMOL^57^ by rotation of helix α2 relative to α1 at the Gln30/Ala31 hinge. For WIPI2d, residues 262-299 of the 6CD region were modeled either (i) as an unstructured loop using SWISS-MODEL^58^ or (ii) with two short helices based on the structure predicted by AlphaFold^59,60^, in two alternative models. Unstructured ATG12 residues 1-52 and WIPI2d residues 1-11 and 362-425 were excluded from the models. The Lys130 side chain of ATG5 was connected to the backbone carbonyl of ATG12 Gly140 by an isopeptide bond. Exposed N- or C-terminal groups at the end(s) of each incomplete structure or truncated construct were neutralized. His183 and His255 at the putative PI(3)P binding sites of WIPI2d were protonated.

All membranes were prepared initially in a coarse-grained representation using the *insane* method^61^ and consisted of 60% DOPC, 20% DOPE, 5% DOPS, 10% POPI, and 5% PI(3)P based on the ER lipid composition^62^. Buckled membranes were constructed using LipidWrapper^63^ by fitting the height (amplitude) of the membrane as a sine function of its *x*-coordinate. For all-atom simulations, the CG2AT2 tool^64^ was used to convert each equilibrated membrane system to an atomistic representation. All simulation systems were solvated with 150 mM of aqueous NaCl, using TIP3P or coarse-grained water. Simulation replicates were independently prepared and equilibrated, with the simulation cells having approximate dimensions of 14 x 14 x 20 nm^3^ for atomistic WIPI2d simulations, 32 x 14 x 28 nm^3^ for simulations of atomistic ATG12–5-L1 on either side of curved membranes (with two copies per replicate system), and 63 x 28 x 38 nm^3^ for coarse-grained curvature-sampling simulations. The *xy* dimensions of buckled membrane systems were fixed during simulation.

Each coarse-grained membrane system was equilibrated for 200 ns, with atomistic systems further equilibrated for 10 ns upon conversion from coarse-grained representation. Harmonic positional restraints were applied to non-hydrogen protein atoms or backbone beads during equilibration, with a force constant of 1000 kJ mol^-1^. In the case of buckled membranes, a weaker (10 kJ mol^-1^) restraint in *z* was also applied to the phosphorus atoms or phosphate beads of lipid headgroups to preserve the initial distance between protein and membrane at the equilibration stage. System temperature and pressure were maintained at 310 K and 1 bar, using the velocity-rescaling thermostat^65^ and a semi-isotropic Parrinello-Rahman barostat^66^ during the production phase. Atomistic and coarse-grained systems were simulated with integration time steps of 2 fs and 20 fs, respectively. For the atomistic simulations, long-range electrostatic interactions were treated using the smooth particle mesh Ewald method^67,68^ with a real-space cut-off of 1 nm, a Fourier spacing of 0.12 nm, and charge interpolation through fourth-order B-splines. The LINCS algorithm was used to constrain covalent bonds involving hydrogen atoms^69^.

Simulation trajectories were analyzed through MDAnalysis 2.0^70,71^ in Python 3.6. The local curvature of buckled membranes during simulation was estimated using the MemCurv software package (https://github.com/bio-phys/MemCurv) following the protocol established by Bhaskara et al. and the same parameter settings as previously described^35^. Sampling at 10 ns intervals, membrane profiles were approximated using a 2D Fourier expansion and optimized by least-squares fitting. The mean curvature *H* at any given position of interest were derived from the shape operator of the approximated profile along the membrane surface^35^.

## Acknowledgements

We thank members of the Aligning Science Across Parkinson’s Team mito911 for advice and discussions. The study is funded by the joint efforts of The Michael J. Fox Foundation for Parkinson’s Research (MJFF) and Aligning Science Across Parkinson’s (ASAP) initiative. MJFF administers the grant ASAP-000350 (to J.H.H., S.M. and G.H.) on behalf of ASAP and itself. The research was also supported by National Institute of General Medical Sciences, NIH, R01 GM111730 (J.H.H.) and the Max Planck Society (S.R. and G.H).

## Figures

**Extended Data Figure 1.**
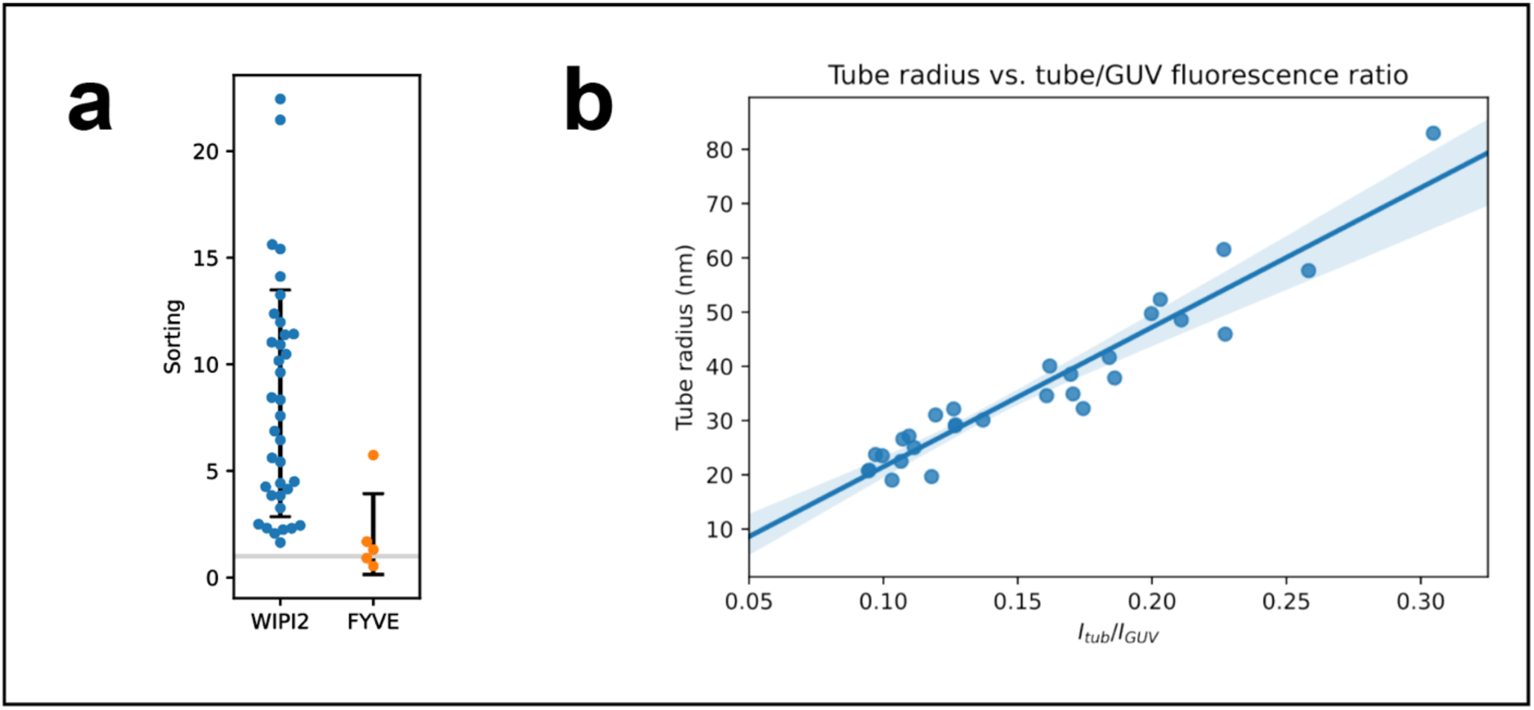
(a) Swarm plot comparison of WIPI2 (blue) compared to FYVE (orange) sorting on membrane tubes. Black bars indicate standard deviation; grey horizontal line at S=1. (b) Tube radius calculated from optical trap force and GUV aspiration pressure vs. ratio of fluorescence intensity of membrane tube to GUV surface. Linear regression plotted as a solid blue line.

**Extended Data Figure 2.**
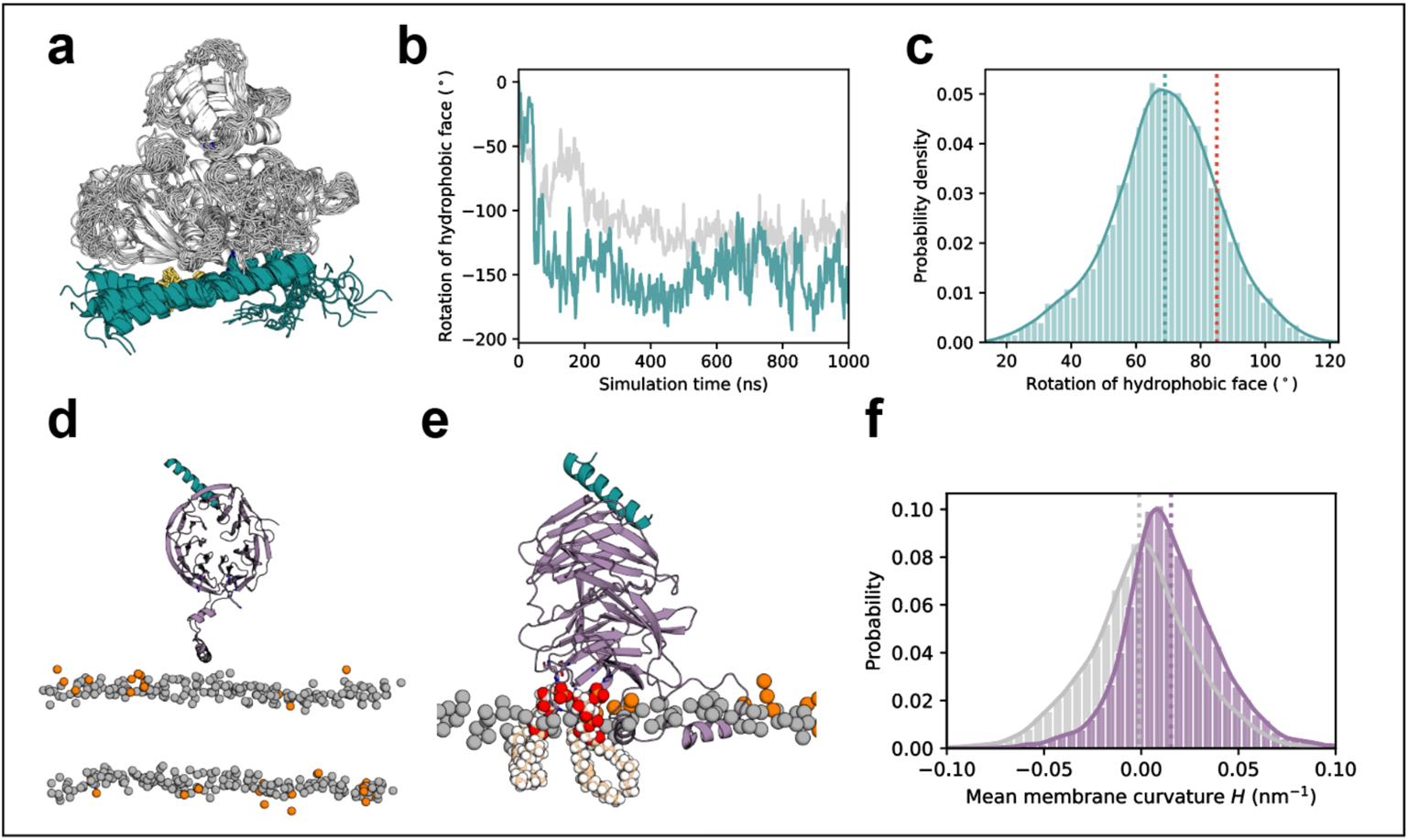
Membrane binding and curvature sensing of ATG12-5–L1 and WIPI2d. (a) Flexibility of ATG16L1 helix 2, illustrated with a superimposition of ATG12-5–L1 conformations sampled at 50 ns intervals during one 1 µs simulation trajectory. The side chains of ATG16L1 helix 1 residues Ile17, Leu21, and Arg24 (cyan) at the interface with ATG5, and of helix 2 residues Phe32, Ile35, and Ile36 of the hydrophobic face (highlighted yellow) are shown in stick representation. (b) Reorientation of the helix 2 hydrophobic face of the aforementioned molecule (cyan) over the course of its simulation, and of a second copy of ATG12-5–L1 (grey) on the opposite membrane leaflet in the same replicate system. At each time point, the rotation angle (relative to the initial conformation) is defined by taking the outward-pointing normal to the plane through the Cα atoms of Phe32, Ile35, and Ile36. Rotation in the clockwise direction, as viewed from the C-terminal end of helix 2 corresponds to a positive value. (c) Distribution of helix 2 hydrophobic face rotation angles in simulations of a remodeled ATG16L1 structure in complex with ATG12-5, over a total simulation time of ~1.8 µs. Rotation angles are measured relative to the conformation in the crystal structure (PDB ID: 4NAW). The remodeled configuration had an initial value approximately in the –*z* direction into the membrane, as indicated by the red dotted line. The mean position of the distribution is indicated by the cyan dotted line. (d) Structure of WIPI2d (magenta) bound to residues 209-230 of ATG16L1 (PDB ID: 7MU2), with the WIPI2d 6CD residues 272-284 and 290-296 modelled as α-helices, placed above a PI(3)P-containing membrane. Residues forming the two putative PI(3)P binding sites in blades 5 and 6 are shown in stick representation. Phosphorus atoms of PI(3)P molecules are shown as orange spheres, with those of all other lipid headgroups colored grey. (e) Snapshot of WIPI2d after forming spontaneous membrane interactions during one (2 µs) simulation replicate, with residues ~271-300 embedded into the membrane. Two PI(3)P molecules, each occupying a binding site on WIPI2d, are highlighted as spheres. (f) Probability histogram of local mean curvature values sampled at the center of mass of WIPI2d 6CD residues 263-300 (magenta) and at random lipid phosphate positions (grey) during the same simulations, with data collected over the final 12 µs of six independent 20 µs replicate systems, each containing two copies of the protein construct, on either side of the buckled membrane.

**Extended Data Figure 3.**
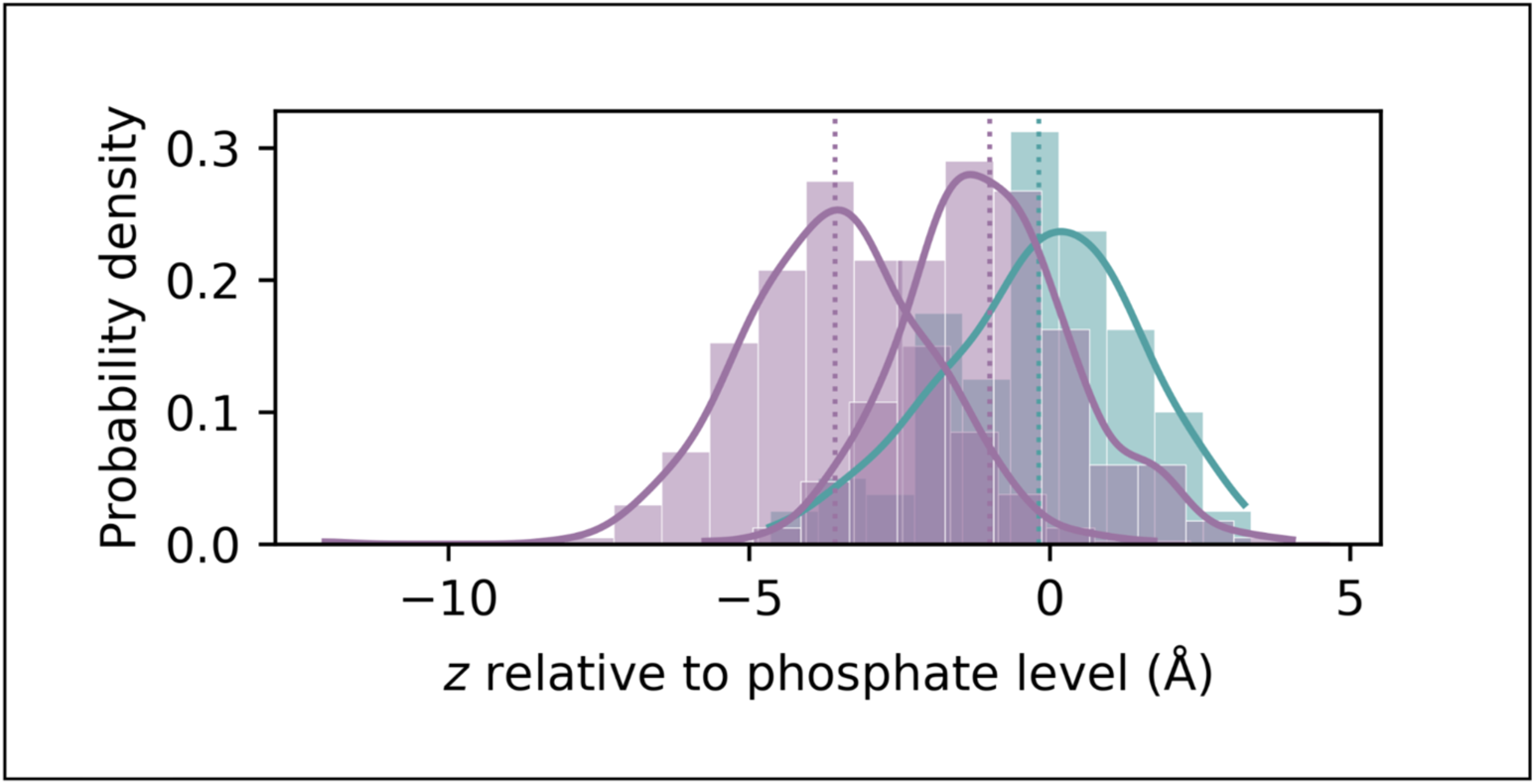
Membrane insertion depths of amphipathic helical elements within WIPI2d (magenta) and ATG16L1 (cyan). The *z*-positions of the center of geometry of WIPI2d residues 273-283 (around a mean value of ~–3.6 Å) and 290-296 (~–1 Å) and of ATG16L1 helix α2 are each measured relative to the level of phosphate groups of the membrane leaflet to which the protein is bound. The distribution of *z*-positions in one representative all-atom simulation replicate of WIPI2d or ATG12–5-16L1 is shown, respectively. Simulation trajectories are sampled at 1 ns intervals after an equilibration period following membrane insertion of the amphipathic helix.

